# Sequence-dependent nucleosome sliding in rotation-coupled and uncoupled modes revealed by molecular simulations

**DOI:** 10.1101/177436

**Authors:** Toru Niina, Giovanni B. Brandani, Cheng Tan, Shoji Takada

## Abstract

While nucleosome positioning on eukaryotic genome play important roles for genetic regulation, molecular mechanisms of nucleosome positioning and sliding along DNA are not well understood. Here we investigated thermally-activated spontaneous nucleosome sliding mechanisms developing and applying a coarse-grained molecular simulation method that incorporates both long-range electrostatic and short-range hydrogen-bond interactions between histone octamer and DNA. The simulations revealed two distinct sliding modes depending on the nucleosomal DNA sequence. A uniform DNA sequence showed frequent sliding with one base pair step in a rotation-coupled manner, akin to screw-like motions. On the contrary, a strong positioning sequence, the so-called 601 sequence, exhibits rare, abrupt transitions of five and ten base pair steps without rotation. Moreover, we evaluated the importance of hydrogen bond interactions on the sliding mode, finding that strong and weak bonds favor respectively the rotation-coupled and -uncoupled sliding movements.

**Author summary:** Nucleosomes are fundamental units of chromatin folding consisting of double-stranded DNA wrapped ∼1.7 times around a histone octamer. By densely populating the eukaryotic genome, nucleosomes enable efficient genome compaction inside the cellular nucleus. However, the portion of DNA occupied by a nucleosome can hardly be accessed by other DNA-binding proteins, obstructing fundamental cellular processes such as DNA replication and transcription. DNA compaction and access by other proteins can simultaneously be achieved via the dynamical repositioning of nucleosomes, which can slide along the DNA sequence. In this study, we developed and used coarse-grained molecular dynamics simulations to reveal the molecular details of nucleosome sliding. We find that the sliding mode is highly dependent on the underlying DNA sequence. Specifically, a sequence with a strong nucleosome positioning signal slides via large jumps by five and ten base pairs, preserving the optimal DNA bending profile. On the other hand, uniform sequences without the positioning signal slide via a screw-like motion of DNA, one base pair at the time. These results show that sequence has a large effect not only on the formation of nucleosomes, but also on the kinetics of repositioning.

## Introduction

Nucleosomes are the fundamental structural unit of eukaryotic chromatin, composed of approximately 147 base pairs (bp) of double stranded DNA wrapped around a histone octamer [1]. Nucleosomes enable genomic DNA to be folded into chromatin and efficiently packed inside the cell nucleus [2]. At the same time, because of the tight association with histones, nucleosomal DNA cannot be usually accessed by other proteins, inhibiting transcription factor association and gene expression [3,4]. Protein binding to a DNA region originally part of a nucleosome usually requires either complete nucleosome disassembly [5] or nucleosome sliding [6] away from the target sequence. The latter mechanism does not involve the complete breakage of histone-DNA contacts.

How are nucleosome positions regulated in the cell? Nucleosome assembly is strongly dependent on the underlying DNA sequence [7], and it has been shown that sequence indeed significantly contributes to the observed pattern of nucleosome positions *in vivo* [8,9]. However, in the complex cell environment many other factors will determine nucleosome positions. For instance, transcription factors will compete with nucleosomes to bind their specific target sites, due to the presence of steric hindrance [4]. Furthermore, many molecular machines, called chromatin remodelers, consume ATP to actively evict nucleosomes or reposition them to new sequence locations [10]. It has also been suggested that remodelers may be particularly important to enhance structural fluctuations, enabling a rapid search of the optimal nucleosome positions [8].

Nucleosomes may also undergo spontaneous repositioning in the absence of active remodelers [11]. While the importance of this mechanism *in vivo* has not been carefully investigated, many *in vitro* studies, e.g. using 2-dimensional electrophoresis [12] and atomic force microscopy [13], confirmed the existence of spontaneous nucleosome sliding. Moreover, experimental and theoretical work suggested that repositioning may occur via a corkscrew motion of DNA [14,15]. However, different repositioning mechanisms that do not involve this kind of rotation-coupled motion of DNA, such as DNA reptation via propagation of loop defects [16], have also been proposed. DNA sequence adds complexity to this problem: Genomes are rich in both positioning and non-positioning sequence motifs that enhance or inhibit nucleosome association [17], and these motifs will likely influence the dynamics of nucleosome repositioning [18].

Here we investigated thermally-activated spontaneous nucleosome sliding dynamics by employing a molecular dynamics (MD) simulation approach. While all-atom MD simulations have been widely used to study the molecular details of nucleosome conformation and histone-DNA interactions [19–22], it would be computationally challenging to reach the time-scales relevant to observe spontaneous sliding. On the other hand, recent studies have shown the effectiveness of coarse-grained (CG) MD simulation in investigating large biomolecular complexes such as nucleosomes [23]. In particular, the combination of the AICG2+ residue-level coarse-grained model for proteins [24,25] with the three-site-per-nucleotide (3SPN) model for DNA [26] proved to be a very successful strategy with many applications to nucleosome dynamics to date [27–29]. Notably, the latest version of the 3SPN model of DNA (3SPN.2C) [30] is designed to reproduce the sequence-dependent DNA flexibility [31], making the model suitable for the study of the influence of DNA sequence on the energetics of nucleosome formation [27].

In this work, we aim to reveal the dynamics of spontaneous nucleosome repositioning using the AICG2+ and 3SPN.2C coarse-grained models, and study the influence of DNA sequence on this process. To achieve this, an appropriate representation of histone-DNA interactions is of particular importance. Firstly, we developed a novel coarse-grained representation of histone-DNA hydrogen bonds, and showed that this potential, together with excluded volume and long-range electrostatics, is necessary to generate stable nucleosomes at low ionic strength and to reproduce the experimental unwrapping behavior observed at higher salt concentrations. Then, we performed MD simulations of nucleosomes using different DNA sequences, and identified two distinct sliding modes: one, coupled to DNA rotation, for uniform and non-positioning sequences, and a second, uncoupled to rotation, for a strong nucle osome positioning sequence. Finally, using a reweighting technique, we investigated the importance of histone-DNA hydrogen bonds in controlling these two sliding modes, finding that weak bonds favor the rotation-uncoupled mode, whereas stronger bonds favor the rotation-coupled one.

## Methods

### Materials

We examined nucleosomes with four DNA sequences of 223 bps; 1) the so-called 601 strong positioning sequence [32] that contains 145 bps (Table 1) [33], flanked by 39-bp linker DNA of polyCG sequence, 2) the polyCG sequence, where one strand has the sequence 5’-CGCG,,,CGC-3’, while the other strand has its complementary sequence, 3) the polyAA sequence made of 5’-AAAA,,, AAA-3’ and its complementary sequence, and 4) a modified polyCG sequence with the addition of two TTAAA positioning motifs at the same locations as those found in the 601 sequence, which we will refer to as the polyCG-601 sequence (Table 1).

**Table 1.**
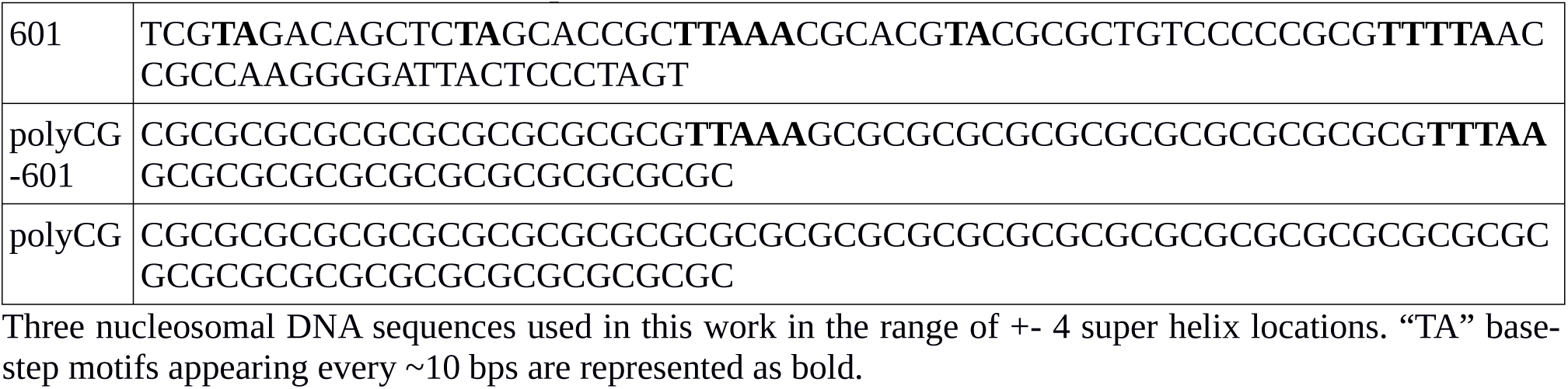
Three nucleosomal DNA sequences

### Reference Structures

For protein modeling, we used the crystal structure with the protein-data-bank id 1KX5 as the reference histone octamer. For the DNA, we used the package 3DNA [34] to generate the reference structures for the 3SPN.2C model to optimally model the sequence-dependent geometric features and flexibility of DNA. To model histone-DNA interactions, we used the 1KX5 and 3LZ0 crystal structures, which are respectively based on the α-satellite [35] and 601 [33] positioning sequences (see section on hydrogen bond interactions for more details).

To prepare the initial structures (Fig 1A), we used the 1KX5 and 3LZ0 crystal structures. For polyCG and polyCG-601 sequences, we used the 1KX5 nucleosome structure that contains 147-bp nucleosomal DNA and added the extra DNA linkers by aligning the last base-pair of an ideal 39-bp segment of DNA to the last base-pair at each end of the nucleosomal DNA, resulting in the addition of two 38-bp DNA linkers at each end, reaching a total of 223 bps of nucleosomal DNA. For the 601 positioning sequence, we used the 3LZ0 nucleosome structure as a template, which has 145 bps instead of 147, and added 39-bp DNA linkers, obtaining the same DNA length of 223 bps. In all the cases, these initial structures are energy minimized using the steepest-descent method before production runs.

**Fig 1.**
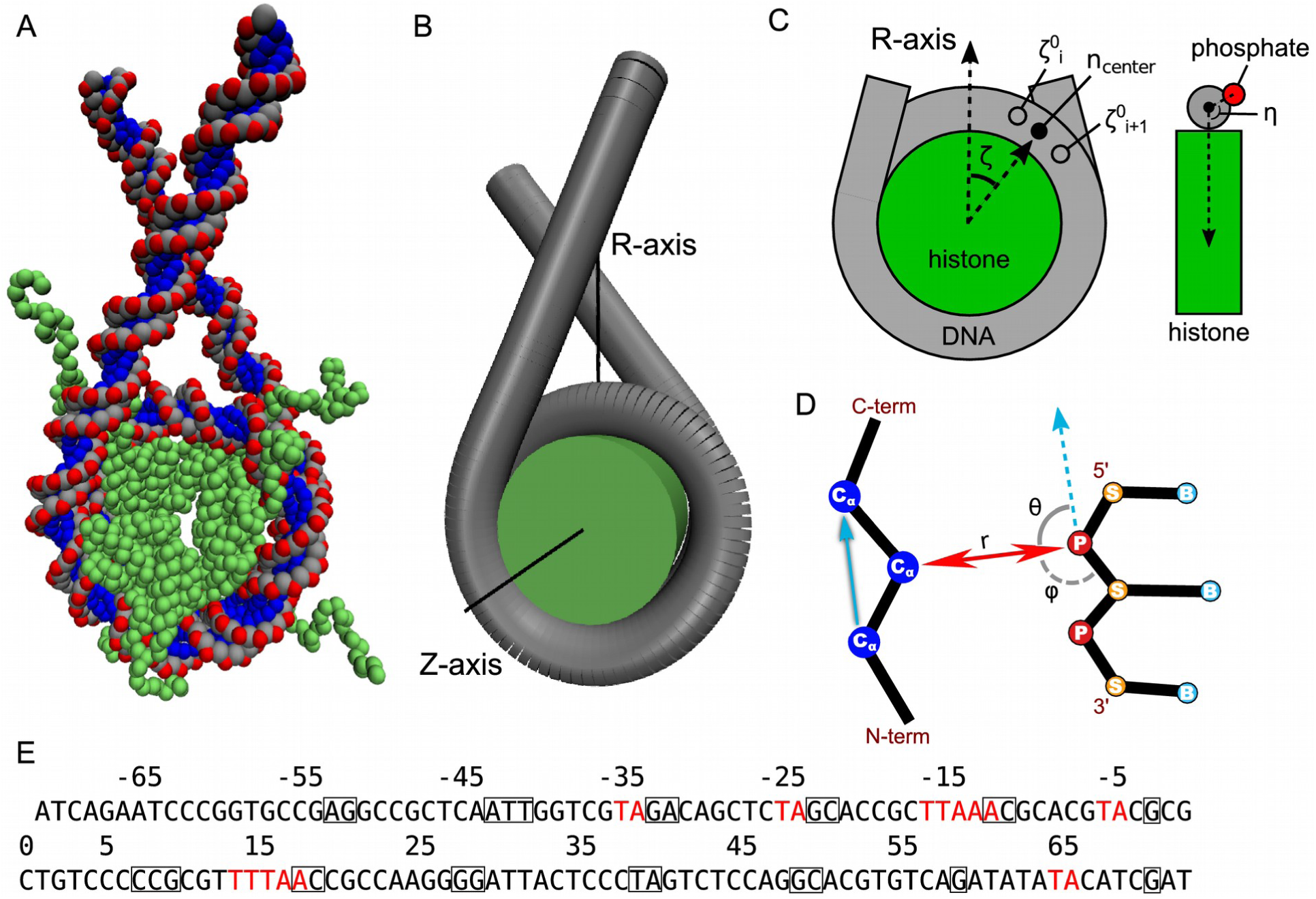
The nucleosome modeling and coordinates. (A) The coarse-grained structure of a nucleosome with linker DNAs. (B, C) The definition of coordinates used in the analysis of the DNA sliding. The η represents DNA rotation coordinate around its long axis, while the ζ is the sliding coordinate, which will be converted to the number of base pairs in the subsequent figures. (D) The coordinates for modeling the hydrogen bond potential: The distance (r) and the two angles (θ and φ). Cα, phosphate, sugar and base beads are respectively indicated by the letters Cα, P, S and B. (E) The Widom 601 DNA sequence. The nucleosome positioning signals are indicated in red. The squared bases represent nucleotides that make hydrogen bonds with histones in the 1KX5 and 3LZ0 crystal structures.

### Protein Modeling

The histone octamer is modeled according to the AICG2+ potential [24,25], where each protein residue is coarse-grained to a single bead located at the corresponding Cα atom. The portion of the histone tails not resolved in the reference 1KX5 crystal structure is modeled using a statistical potential that reproduces the residue-dependent probability distribution of angles and dihedral angles between consecutive residues as observed in a database loop crystal structures [36].

### DNA Modeling

To model the double-stranded DNA we employed the sequence-dependent 3SPN.2C coarse-grained model [30]. Within this model, each nucleotide is represented by three beads corresponding to sugar, phosphate and base groups. The model has been parameterized by matching the experimental DNA melting temperature, persistence length, and average base-pair and base-step parameters for all the ten unique base-step types [30,31]. The accurate representation of sequence-dependent effects is of particular importance for our investigation, and this model has already been shown to reproduce well the experimental dependence of nucleosome formation on the underlying DNA sequence [27].

### Protein-DNA Interaction

The histone octamer interacts with the DNA via excluded volume, Debye-Huckel electrostatics and a novel coarse-grained potential representing hydrogen bonds, which we develop here. The excluded volume is modeled by a r^12^ repulsive potential, where the particle radii are bead-type dependent and they have been estimated from the minimum distances between each pair of bead types observed in a database of protein-protein and protein-DNA complexes [37]. With respect to the original parameters in Ref. [37], the radii have been uniformly rescaled by a factor of 1.1, which prevents the histone tails from being able to insert between two DNA strands. To represent long-range electrostatic interactions between histone proteins and DNA, and within the DNA, we used the Debye-Huckel approximation with a temperature- and salt concentration-dependent dielectric constant, as described in Ref. [26]. For DNA-DNA electrostatic interactions, following the recommended settings [26,30], we set a charge of −0.6e on each phosphate bead, which takes into account the Oosawa-Manning condensation of counter ions around DNA and it results in the correct DNA persistence length. On the other hand, for protein-DNA electrostatics, we set the phosphate charges to −1e as in Refs. [37] and [27], whereas the charges on the globular part of the histone octamer have been estimated using the RESPAC method [38]. In RESPAC, the coarse-grained charges are optimized so that the resulting electrostatic potential provides the best approximation to the all-atom electrostatic potential of the protein in the native reference 1KX5 crystal structure. The optimization procedure has been performed at 100 mM salt concentration, but the resulting charges show very low sensitivity to this particular value, so that the same set of charges can be used to run simulations at ionic strengths used in this work. The RESPAC method is only appropriate where the protein remains close to the reference native structure during the MD simulation, therefore for the flexible histone tails (up to the first structured alpha helix in the histone) we employed the standard residue unit charges: +1e for lysine and arginine, and −1e for aspartic and glutamic acids.

### Hydrogen Bond Interactions

Many, but not all, coarse-grained simulations of nucleosomes reported so far employed Go-like potentials for the histone-DNA interaction to ensure that the nucleosome core structure is stable and close to the observed crystal structure. While this approach is convenient for the study of many important processes such as nucleosome breathing [28] and assembly [39], it cannot be applied to the study of nucleosome repositioning, since these potentials assume specific interactions at the prefixed positioning and thus are not invariant under DNA sliding with respect to the histone core. To overcome this limitation of standard Go potentials, we follow a similar approach to the one developed by Tan et al. (Tan and Takada, in preparation) to model the specific sequence-dependent binding of transcription factors to DNA. In the current study, we introduce this specific potential as the histone-DNA hydrogen bonds that stabilize the nucleosome structure. These hydrogen bonds are formed between a set of histone residues and the DNA backbone phosphates located at the half-integer super-helical locations, where the DNA minor groove faces the octamer. We set this potential to be invariant under a rotation-coupled repositioning of the DNA, since every phosphate bound to the protein will be simply replaced by a new phosphate with the same relative orientation.

To achieve this, we could in principle create the same Go-like contact between each protein acceptor and every single phosphate in the DNA. However, the problem with this strategy is that the standard 12-10 Lennard-Jones potential normally employed is too broad, and each hydrogen bond will be counted multiple times, not only between the protein residue and the correct phosphate, but also including other neighboring phosphates.

This issue can be overcome by making the potential highly specific, using both distance- and angle-dependence to represent the formation of a hydrogen bond (HB) (Fig 1D), of which idea came from recent coarse-grained DNA modeling [26]. In our model, we define the contribution to the potential energy from these bonds to be:

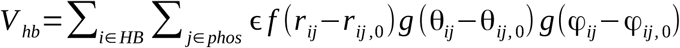

where the first sum runs over the list of native HBs identified in the reference nucleosome crystal structures and the second sum runs over all the DNA phosphate beads, ensuring the invariance of the potential under a rotation-coupled repositioning of DNA. ε is the energy parameter that controls the hydrogen bond strength, *r*_*ij*_ is the distance between the Cα bead of the *i*-th HB-forming residue and the *j*-th phosphate, *θ*_*ij*_ is the angle between the vector connecting the *i*-th HB-forming residue to the phosphate and the vector connecting the two residues neighboring the bond-forming one along the polypeptide chain (see Fig 1D), *φ*_*ij*_ is the angle between the HB-forming residue, the considered phosphate and the sugar bead in the same nucleotide of the phosphate. The parameters with the subscript 0 are the corresponding distance and angles of the considered HB *i* found in the native structure, which are used as a reference to evaluate the formation of each specific bond. The functions *f* and *g* control the distance- and angle-dependence of the potential, which take a value close to 1 where the argument is close to 0, i.e. when the distance or angle variable observed during MD is close to the reference, and decreases as the argument deviates from zero. Their precise functional forms are given by:

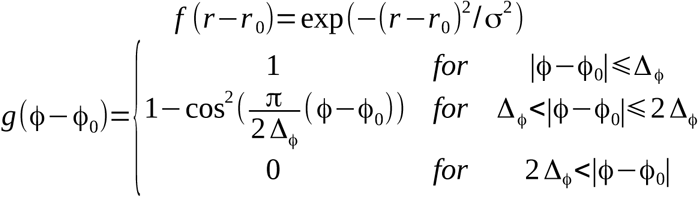

Where the potential widths σ and Δ_φ_, within which a bond is considered to be formed, are respectively set to 1 Å and 10 degrees. These values have been identified by requiring the HBs in the reference 1KX5 crystal structure to be well defined to their specific phosphate groups, without including favorable energetic contributions coming from the neighboring phosphates, which would amount to a double-counting of the interactions and would occur if the widths are too large. The energy constant ε was set to 1.2 *k*_B_*T,* which is about the smallest value required to stabilize the nucleosome structure against large deviations from the reference crystal, while still allowing the expected nucleosome disassembly at large salt concentration (see model validation). The list of HBs and their reference distance and angle parameters have been generated from the HBs found in any of the nucleosome crystal structures with PDB id 1KX5 and 3LZ0 using the software MDAnalysis [40] with the default settings. Flexible tails are excluded from the analysis because in the potential that evaluates the formation of a hydrogen bond we are assuming a stable near-native protein conformation (the bonds formed by the tails only account to a small portion of the total histone-DNA hydrogen bonds). The reference distance and angle values of each bond are obtained from an average over the two crystal structures and the two symmetric halves of the nucleosome for each structure.

### Molecular Dynamics Simulations

All the CG simulations were performed by the CafeMol package version 3 [41]. The simulations were conducted by Langevin dynamics with default parameters at a temperature of 300 K. During the simulations of the unwrapping analysis, the ionic strength was varied from 100 to 1000 mM of mono-valent ions, and for each ionic strength, 10 independent 10^8^ MD-step simulations were carried out. For the production runs, all the system is placed into a sphere with radius 80 nm and repulsive walls. The ionic strength was set to 200 mM, and we carried 100 independent 10^8^ MD-step simulations for each considered sequence (polyCG, polyCG-601, and 601).

### Analysis

To analyze the mode of nucleosome repositioning, we considered two angular coordinates: the sliding coordinate ζ and the DNA rotation coordinate η (Fig 1B, C). The former is defined by the angle between the vector from the histone core center of geometry to the center of the base pair initially at the dyad and the vector corresponding to the nucleosome symmetry axis; whereas the latter, η, is defined by the angle between the vector from the DNA axis to the 1st-strand phosphate of the base pair initially at the dyad and the vector from the DNA axis at the same base pair initially at the dyad to the histone core center of geometry. To extract the position of the DNA axis at each base pair, we firstly define 10 spline lines connecting each phosphate group of residue i in the first DNA strand with the phosphates of residues i-10 and i+10 (and i-20, i+20 and so on), representing contours on the DNA tube. Then, for each 1 st-strand phosphate group, we compute the closest points on each of the 10 contours amongst the set of points obtained by subdividing each spline segment into 10 equal parts. Finally, for each base pair corresponding to the phosphate group, we define the DNA axis position as the center of the circle obtained from a fit of these 10 closest points. The nucleosome symmetry axis has been obtained by fitting a straight line to the set of the centers of geometry of the symmetric pairs of Cα beads in the histones (e.g. the center of geometry of the Cα beads with residue id 51 in the 1st and 2nd H3 histones, and similarly for other symmetric residue pairs; flexible tails were not taken into account).

In order to simplify the understanding of the sliding dynamics, we convert the unit of the sliding coordinate ζ from angles to number of base pairs. To do this, we first make a table that maps base pair indexes to ζ angles as obtained from the initial nucleosome configuration. For each snapshot in the trajectories, from the table, we then find the two neighboring nucleotides closest to the obtained ζ angle. Then we calculated the number of slid base pairs corresponding to ζ via linear interpolation between the two base pair indexes.

## Results

### Model validation 1: Nucleosome stability of a positioning sequence

The highly-bent nucleosomal DNA is stabilized by a strong electrostatic attraction with histones and by more local interactions primarily via a network of hydrogen bonds to the histone octamer core residues [42]. Since we cannot employ the Go-like potential for the histone-DNA interactions in this study, an alternative and accurate modeling for the histone-DNA interactions is indispensable. Here, we test our CG modeling on nucleosomes formed with the 601 positioning sequence [32]. This sequence is well-known for the presence of several TA base-step positioning motifs (see Table 1) [8] which prefer to localize at nucleosome regions where the DNA minor groove faces the histone octamer, due to the intrinsic bending of DNA [27].

We began with a simple CG model where only the electrostatic interactions are included as the attraction between histones and DNA [27]. We performed CGMD simulations using the 601 sequence flanked by 39-bp polyCG sequences in both termini at the salt concentration of monovalent ions 200mM. With this condition, the nucleosome is stable experimentally. In the initial structure, the 601 sequence is wrapped by the histone octamer while the 39-bp polyCG segments form the linker DNAs. The resulting root-mean-square-deviation (RMSD) from the reference structure 1KX5 is plotted as a function of simulation timestep in Fig 2A for three representative trajectories (blue, red, and green curves). The RMSD was calculated for the central part of DNA (the segment that is initially located between −1 and +1 super helical locations) after alignment of the globular part of the histone octamer. First, we see that the native-like state with the averaged RMSD of ∼ 0.5 nm is only marginally stable (only the green trajectory stayed in this state for significant time) (see Fig 2C). Instead, all the three trajectories stayed much longer time with the averaged RMSD of ∼1.5 nm. In this state, the nucleosomal DNA is still well-wrapped around the histone core, while DNA is slid by ∼5 bps so that the major groove positioned around the sites where the minor groove exists in the initial structure, likely representing an artifact due to inaccurate histone-DNA interactions (see Fig 2D). On top, occasionally, nucleosomal DNA went too far away from its favorable position (Fig 2E). Thus, we conclude that the long-range electrostatic attractive interaction alone is not specific enough to stabilize the nucleosomal DNA at high precision, even though the DNA can be well-wrapped around the histone core.

**Fig 2.**
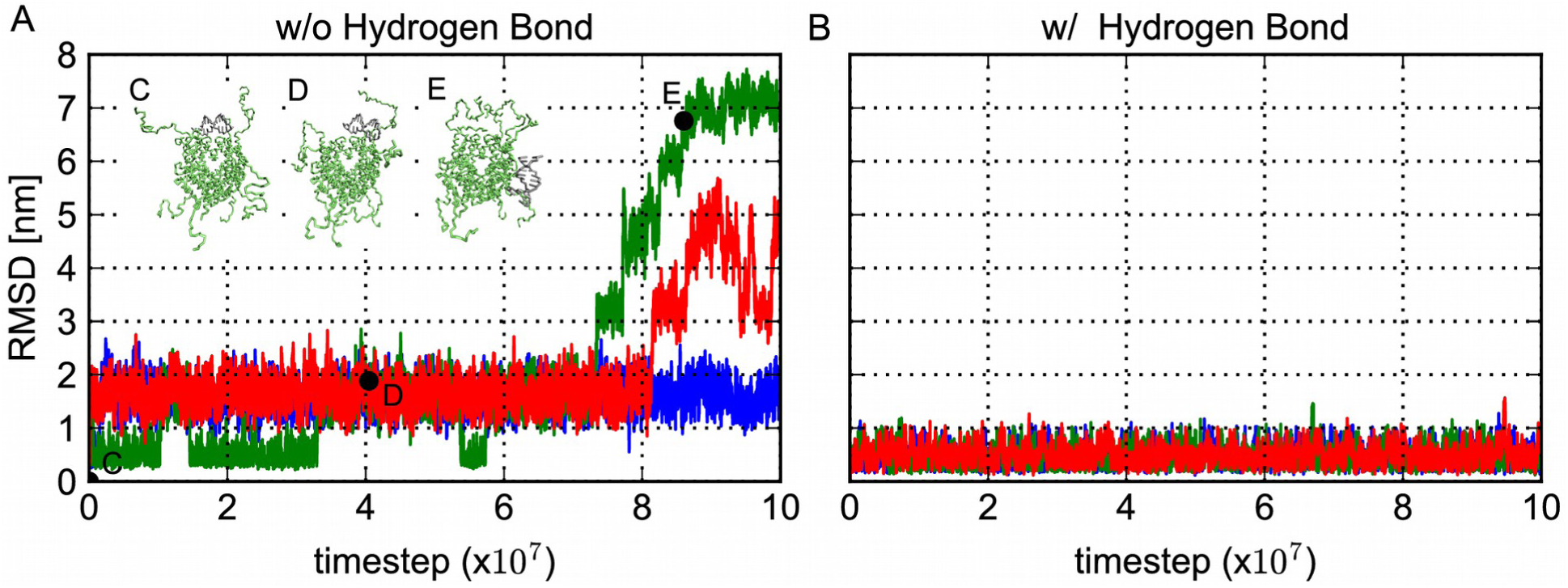
The representative time course of the root mean squared deviation (RMSD) of the nucleosomal DNA. Three representative time course of the RMSD of the nucleosomal DNA from the reference structure 3LZ0 for the Widom 601 sequence and snapshot of the structure corresponding to time points. (A) Three representative time courses of RMSD of the central 20 base pairs of DNA around the dyad when structures were superimposed on the histone octamer core, using a model without histone-DNA hydrogen bond interactions. (B) Three representative time courses of RMSD after the addition of histone-DNA hydrogen bonds. The RMSD was calculated using the same DNA regions described above.. (C-E) Snapshots of structures at the points indicated as black circles in A. Only the histone octamer and the DNA region used to calculate the RMSD are depicted. Structure C is close to the initial structure and D corresponds to the dominant 5-bp-shifted configuration appeared in the simulation without the hydrogen bond potential.

Next, in order to improve the representation of the system, we added the hydrogen bond (HB) potential, as well as the electrostatic interactions, to the histone-DNA interactions. The HB potential depends not only on the distance between the amino acid and the phosphate group of the interacting pairs, but also on the two related angles (see Methods), making the potential highly specific. For the same DNA sequence and the initial structure as above, we performed CGMD simulations with the HB potential. The representative RMSD time courses depicted in Fig 2B show stable fluctuations around ∼ 0.5 nm throughout the simulation time. The nucleosomal DNA resides in the positions well close to those in the crystal structures. These results suggest that combination of the HB potential and the long-range electrostatic interaction is sufficient to stabilize the nucleosomal DNA at high precision in the current CG modeling.

Notably, in the above simulations, we utilized the HB strength parameter ε =1.2 *k*_B_*T*. In some preliminary tests, we found that the ε smaller than 1.0 *k*_B_*T* lead to transitions to ∼5-bp-shifted state. Instead, ε larger or equal to 1.2 *k*_B_*T* resulted in small fluctuations around the crystal structure.

### Model validation 2: nucleosome unwrapping depending on the salt concentration

As the second test of the histone-DNA interactions in our CG molecular model, we examined the dependence of nucleosome unwrapping stability on the ionic strength. We performed MD simulations for the three DNA sequences of 223 bps; the 601 positioning sequence, the polyCG sequence, and the polyAA sequence. Notably, the polyAA sequence is well known to have an inhibitory effect to form nucleosomes. We prepared the initial structures where the central 147 bps are wrapped by the histone octamer. For each DNA sequence and the varying salt concentration between 100 and 1000mM, we produced 10 independent MD simulations of 10^8^ MD steps with different stochastic forces. To examine the convergence to the equilibrium, we also performed CG simulations from fully unwrapped DNA structures with the pre-formed histone octamer: we note that, in reality, the histone octamer is known to be unstable under physiological conditions without the nucleosomal DNA and thus, the initial structure used here does not represent the realistic state. The results indicated that the structural ensemble from the fully-unwrapped and fully-wrapped structures is nearly the same after about 2x10^7^ MD steps. Thus, in the production run of 10^8^ MD steps, the first 2x10^7^ MD steps were discarded.

Fig 3 plots the resulting salt-concentration dependent DNA unwrapping from the histone core. At each salt concentration the average number of DNA phosphate groups within 1.2 nm from the globular part of the histone core was numerated. These titration curves show standard sigmoidal shapes, in which the critical salt concentration that causes significant DNA unwrapping from the histone core is around 500 mM. This is consistent with the range between the ionic concentrations reported by experimental studies for the 601 sequence [43,44]. We performed additional simulations using the 601 sequence with several hydrogen-bond interaction strengths at the high salt concentration of 1000 mM. The average number of histone-DNA contacts at the end of long 10^8^ steps simulations is plotted on the inset of Fig 3. These results show that complete nucleosome disassembly occurs for a wide range of hydrogen bond strengths, and only for ε greater than 2.4 *k*_B_*T* the nucleosome becomes clearly overstabilized. Thus, for the nucleosome with the 601 sequence being stable near the crystal structure at 200 mM and being disassembled at 1000 mM, the acceptable range of hydrogen-bond strengths may roughly be 1.2 *k*_B_*T* ≤ ε ≤ 2.4 *k*_B_*T*.

**Fig 3.**
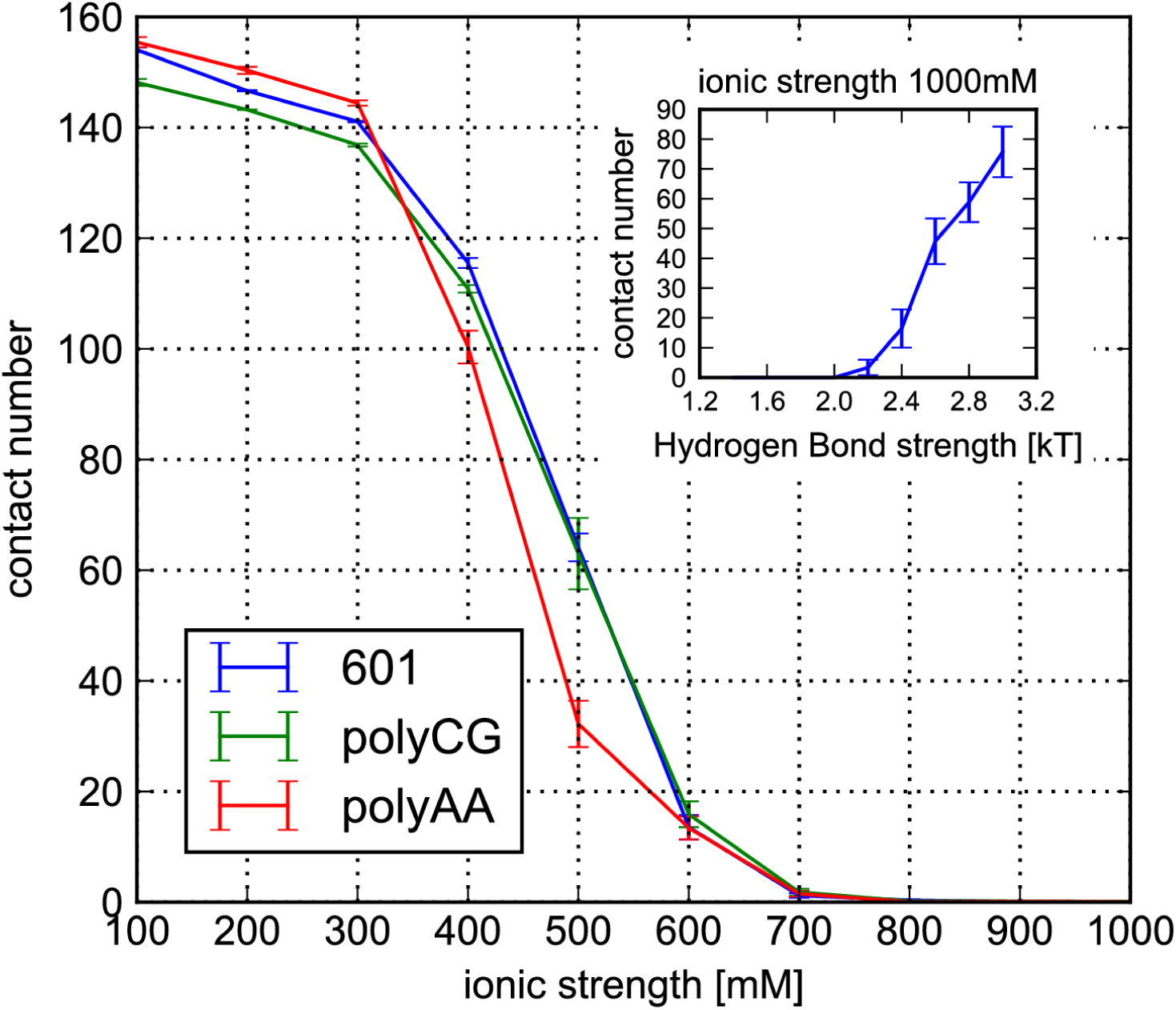
DNA unwrapping as a function of ionic strength for the three DNA sequences. The average number of contacts formed between phosphate groups and histone octamer (excluding tails) are plotted for the hydrogen bond strength ε = 1.2 *k*_B_*T*. The standard errors are plotted as error bars. (inset) The average number of contacts obtained from simulations at ionic strength of 1000 mM as a function of the hydrogen bond strength.

Notably, the 601 and polyCG sequences show a similar behavior in the unwrapping simulation, allowing us to compare the spontaneous sliding of polyCG and 601 at same ionic strength. In contrast, the polyAA sequence is less stable, which is in harmony with the well-known inhibitory effect of the polyAA sequence to form nucleosomes.

### Sequence-dependent nucleosome sliding

In order to observe thermally-activated nucleosome sliding, we performed larger-scale MD simulations of nucleosomes with linker DNAs. To investigate the DNA sequence dependence of the sliding, we use the three distinct DNA sequences of 223 bps (Table 1); the 601 positioning sequence (the same as above), a 2-bp periodic polyCG sequence (the same as above) as a representative of uniform non-positioning sequences, and a modified polyCG sequence with the addition of two TTAAA positioning motifs at the same locations as those found in 601 (polyCG-601 sequence). In order to allow space for sliding, polyCG and polyCG-601 sequences had 38 bps of polyCG linkers flanking each side of the central 147 bps of nucleosomal DNA; similarly, the 601 sequence had 39 bps of polyCG linkers and 145 bps of nucleosomal DNA (as found in the 3LZ0 crystal structure). We expected the polyCG sequence to be one of the most mobile sequences because the nucleosome energy landscape will be invariant under a DNA shift by 2 bps, as opposed to the 601 sequence, which has several AT base-pair step positioning signals placed roughly every 10 base pairs where the minor groove sharply bends towards the histone octamer. The polyCG-601 sequence is expected to display an intermediate behavior between the two. We performed 100 independent MD simulations of 10^8^ MD steps at the salt concentration of monovalent ion 200 mM. To quantify the sliding, we defined the sliding coordinate as the angle ζ between the nucleosome symmetry axis and the vector from the histone core center of geometry to the base pair center (see Fig 1B, C). For clarity, we express the sliding coordinate in base pairs, mapping the angle ζ to the number of base pairs by using the initial configuration (see Methods for details).

Fig 4A plots, for the polyCG sequence, three representative time courses of the central nucleotide position, which was initially placed at the bp index zero. We find frequent and reversible transitions, which apparently suggest one-dimensional random-walk with the step size of one bp. We also plotted the residence probability of the central nucleotide during the entire time course (Fig 4D). Since the simulation started with the bp index zero, the probability distribution has a peak at zero bps and takes a Gaussian shape.

**Fig 4.**
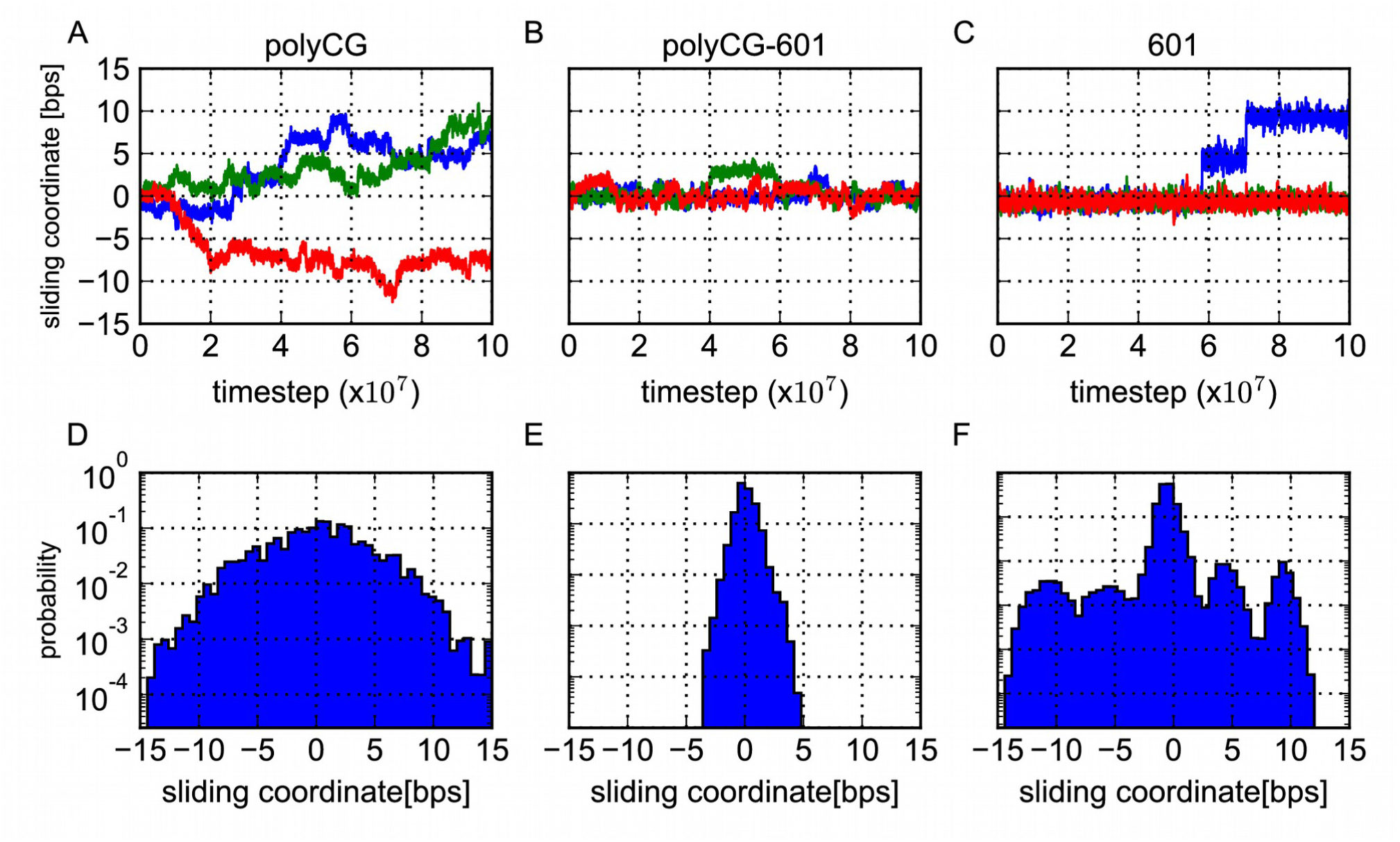
Resprentative time courses and statistical distributions of the sliding coordinate. Representative time courses (A-C) and statistical distributions (D-F) of the sliding coordinate (the deviation of the base pair initially at the dyad from its starting configuration) for each considered sequence: polyCG (A,D), polyCG-601 (B,E) and 601 (C,F). The time courses were moving averaged over 10 time points.

The polyCG-601 sequence also shows frequent repositioning events, but it is significantly more stable than the polyCG sequence (Fig 4B). Notably, the bp index often comes back to the initial position, suggesting that the sliding is limited up to ±4 bps. This is due to the TTAAA motifs optimally localized at the regions of high minor groove bending. The residence probability highlights this point, exhibiting a narrower and bound distribution.

In contrast to the former two sequences, the strong positioning 601 sequence shows nearly no sliding during the entire time course for many trajectories. Only in 14 out of 100 trajectories, however, we found abrupt transitions of ∼5 bps. Once the 5-bp transitions occur, we often observe a subsequent 5-bp transition either to the same or to the opposite directions (see the blue trajectory in Fig 4C). In the 100 trajectories, we obtained 7 events to reach ±10-bp states. The residence probability distribution in Fig 4F clearly shows five peaks/modes. We note that, even though 5-bp-shifted state and 10-bp-shifted state have nearly the same probabilities, a close analysis suggests differently. Since the 5-bp-shifted state is next to the zero bps initial position, this state was visited more times than the 10-bp-shifted state, while the lifetime of 5-bp-shifted state was shorter. This clearly indicates that, in terms of the free energy, the 10-bp-shifted state is more stable than the high-energy intermediate of 5-bp-shifted state.

In summary, the repositioning behavior seems to be different among the three examined sequences. While the polyCG sequence slides frequently with the step size of one bp, the 601 sequence slides much rarely with a minor step size of 5 bps and a major step size of 10 bps. To our surprise, the sliding of the polyCG-601 sequence is not “in between” the polyCG and the 601: within the same simulation time, the sliding was the most restricted for the polyCG-601 sequence. Next, we analyze the DNA sliding motions in more detail.

### Two distinct sliding modes

We analyzed the motion of DNA in detail from the configuration of the DNA base pair initially located at the dyad relative to the nucleosome core. We considered two coordinates; the sliding coordinate and the rotation coordinate. The former is defined in the previous section whereas the rotation coordinate is defined by the angle between the vector from the axis of DNA to the phosphate of the base pair initially at the dyad and the vector from the axis of DNA to the histone core centroid (Fig 1C). If the nucleosomal DNA slides around the histone core in the screw-like manner, changes in the two coordinates would be coupled, otherwise not. In the cases of polyCG and polyCG-601 intermediate sequence, the time courses of the two coordinates are fully coupled (Fig 5A, B). On the contrary, in the case of the sudden jumps observed with the 601 sequence, the time courses of DNA sliding and rotation did not show any coupling behavior, with the DNA orientation remaining constant over time (Fig 5C, F). These results show that there are two distinct sliding modes, a rotation-coupled mode for polyCG and polyCG-601, and a rotation-uncoupled mode for 601.

**Fig 5.**
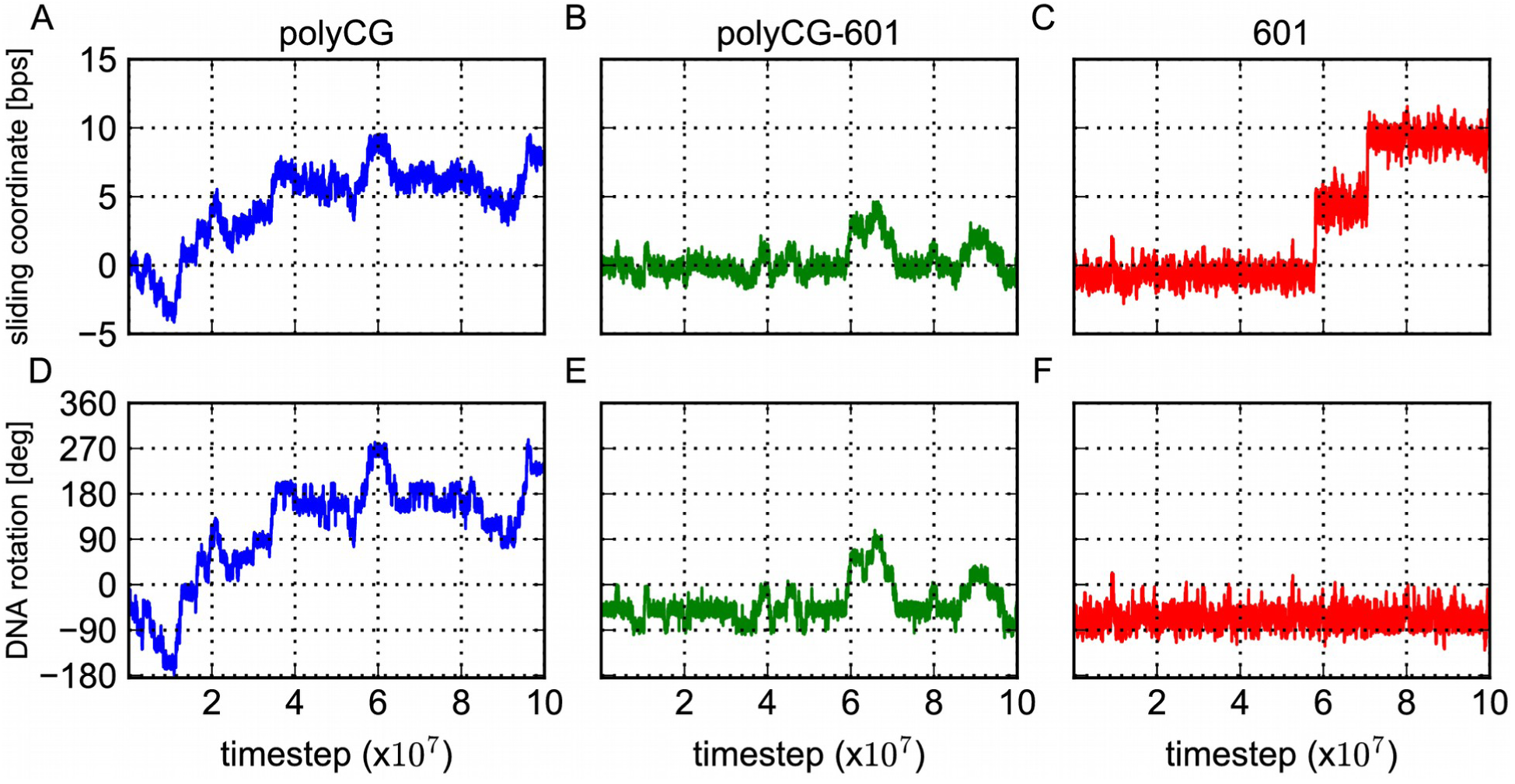
Representative time trajectories of the sliding and DNA rotation coordinates. Representative time trajectories of the sliding (A-C) and DNA rotation (D-F) coordinates for the 3 distinct sequences: polyCG (A,D), polyCG-601 (B,E) and 601 (C,F). The coupling between sliding and rotation signals the screw-like movement of DNA (as in polyCG and polyCG-601), whereas the absence of coupling indicates a jump-like movement (as in 601).

To show the coupling more clearly, we plotted the logarithm of the probability distribution in the two dimensions; the rotation and sliding coordinates (Fig 6). In the case of polyCG, we see a series of high probability spots corresponding to every single bp sliding (Fig 6A). Interestingly, the high probability basins lie along a diagonal line with a slope of 360 degree rotation per ∼10 bps sliding, corresponding to the DNA helical pitch. This signals a perfect rotation-coupled sliding of DNA that enables repositioning without inducing any distortion to the overall nucleosome conformation (Fig 6D, E). The polyCG-601 sequence displays a similar probability distribution to that of polyCG, with the difference that the introduction of the TTAAA motifs breaks the rotational invariance, still allowing the rotation-coupled motion but biasing it towards the initial optimal configuration (Fig 6B, F, G, H).

**Fig 6.**
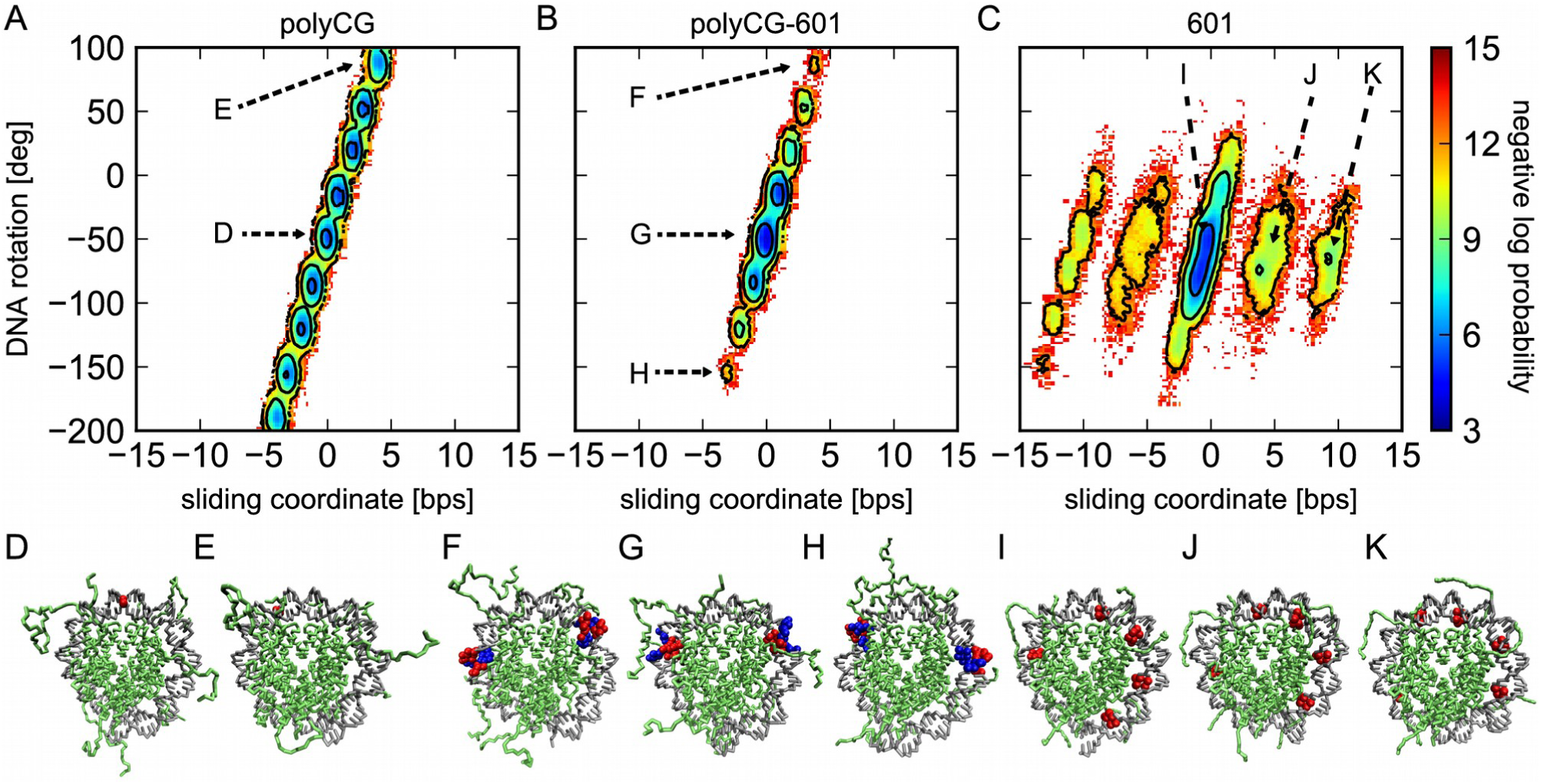
Log-probability of the sliding and rotation coordinates. The colors represents the negative log-probability weighted by *k*_B_*T* in kcal/mol. (A) The polyCG sequence shows that most conformations lie along a straight line having a slope of approximately 360 degree over 10 bps, indicating the ideal screw-like rotation of DNA. (B) The polyCG-601 histogram also shows a straight line but the diffusion is now more limited. (C) For the 601 sequence we find 5 distinct basins corresponding to conformations shifted by 5 bps relative to each other. Within each basin, small screw-like rotation of DNA is also observed. (D-K) Some snapshots corresponding to states pointed in A, B, and C.

In the case of the 601 sequence, we find two distinct motions. First, we see five isolated islands, corresponding to −10, −5, 0, +5, and +10-bp-shifted states, which all have similar rotation coordinate values. Thus, the transitions among the 5 states are not accompanied by the DNA rotation. After sliding by 10 bps the nucleosome conformation will be similar to the optimal initial state, with TA base pair steps located at the inward-bending minor groove regions. Secondly, within each island, we find the rotation-coupled sliding up to ± 2 bps, which is qualitatively similar to that observed for polyCG and polyCG-601 cases.

### Free energy estimates with varying hydrogen bond strengths

From the free energy profiles in Fig 6, we identify five metastable states for the 601 nucleosome, corresponding to −10, −5, 0, 5, and 10 bps repositioning from the initial configuration. For shifts by 5 bps, the configuration looks rather different from the standard nucleosome crystal structures: here the minor groove locations correspond to what was initially occupied by the major grooves. In this intermediate state, long-range electrostatic interactions are not significantly affected; however, most hydrogen bonds are broken. In order to investigate how hydrogen bond interactions influence the rotation-uncoupled repositioning mode via this intermediate state, we performed the reweighting estimate of free energies along the sliding coordinate using different hydrogen bond strengths.

It should be noted that, in this analysis, we restrict ourselves to two states only, the 0-bp and +5-bp-shifted states, because of the limited sampling. To estimate the relaxation time scale within the two states, we performed additional 50 MD simulations starting from +5-bp-shifted state, and considering the 27 trajectories that came back toward the initial configuration we estimated the time scale for the transition from the +5-bp-shifted to the 0-bp state being 1.46x10^7^ MD steps. Since the typical time for a transition from the 0 to the +5-bp-shifted state is orders of magnitude longer, the relaxation time of the whole system is well approximated by the former and shorter time scale. Thus, in the ensemble of the 100 independent MD simulations, we discarded the first 3x10^7^ MD steps to remove the initial-configuration bias.

The resulting free energy profiles in Fig 7A show that increasing the hydrogen bond strength drastically affects the relative populations of 0-bp and +5-bp-shifted states in the rotation-uncoupled sliding mode: with a HB strength of 1.5 *k*_B_*T* (only 25% higher than the setting used in our MD), the free energy difference reaches 10 *k*_B_*T*, strongly inhibiting the rotation-uncoupled sliding mode.

**Fig 7.**
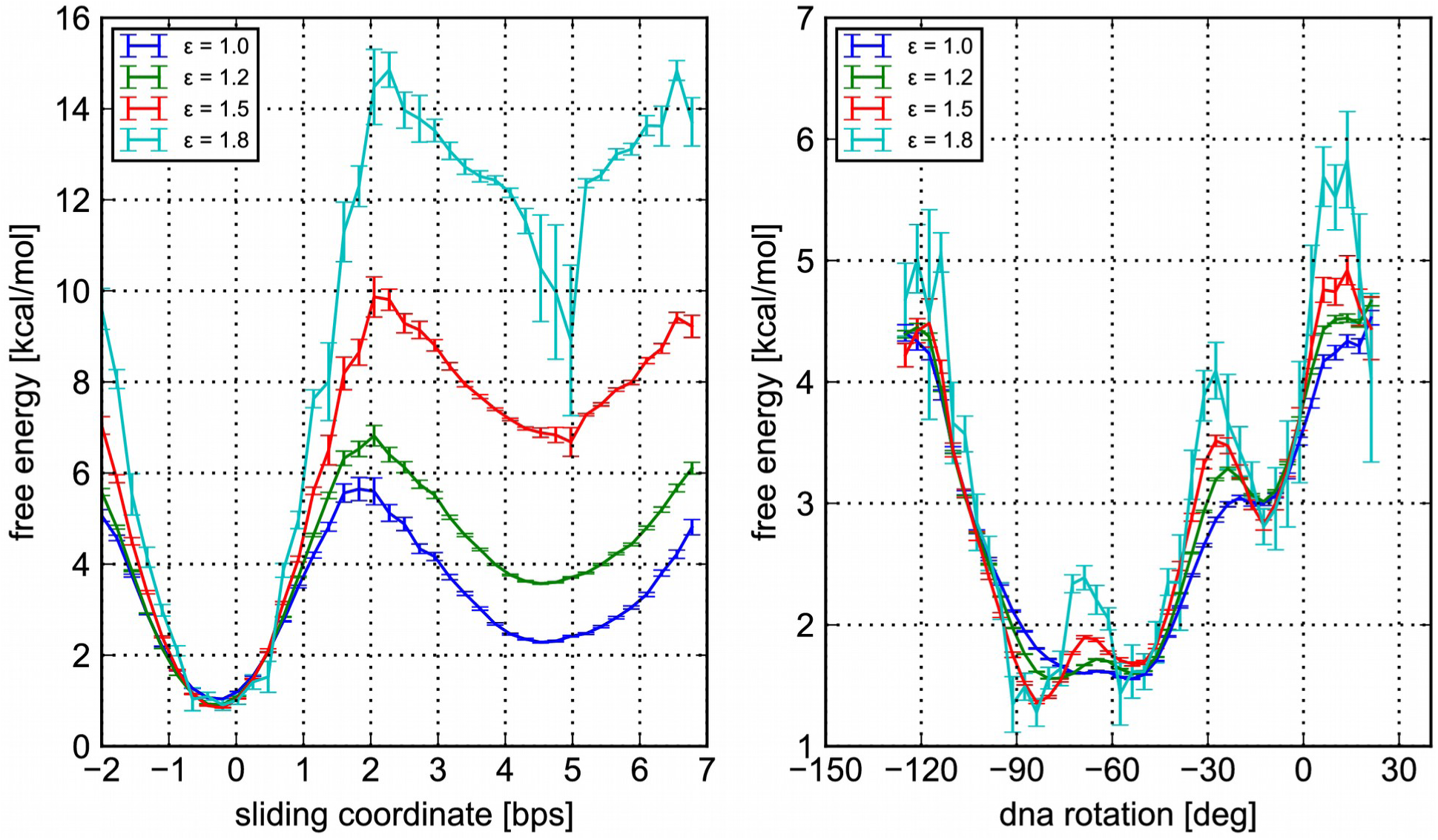
Free energy profiles of the 601 sequence along the sliding and rotation coordinates for different hydrogen bond strengths ε. For ε equal to 1.0, 1.5 and 1.8 *k*_B_*T*, the profiles have been obtained by reweighting the configurations observed during the simulations with ε = 1.2 *k*_B_*T*. (A) Free energy curves along the sliding coordinate. (B) Free energy curves along the DNA rotation coordinate.

For comparison, we also estimated the free energy profile of the 601 sequence within the central island along the rotation-coupled sliding up to ±1-bp-shifted states (Fig 7B). The results show that a change in the HB strength does not significantly affect the free energy profile for the rotation-coupled repositioning mode of the 601 sequence. This is reasonable because the rotation-coupled repositioning does not break the HB except at transient states.

Put together, these results suggest that for stronger hydrogen bonds, the screw-like rotation-coupled mode may become the main mode of thermally-activated spontaneous sliding of nucleosomes, even with strong positioning sequences such as 601.

## Discussion

The sequence dependency of the dominant sliding mode is a result of a balance between DNA’s local bending energy and hydrogen bond breaking penalty. The rotation-coupled motion of DNA does not affect histone-DNA hydrogen bonds, which have to be broken only temporarily when base pairs are exchanged, but it changes the bending profile of the DNA with a periodicity of about 10 base pairs. On the other hand, rotation-uncoupled sliding does not affect DNA bending, but it proceeds via a long-lived intermediate state where most hydrogen bonds are broken.

For a uniform sequence such as polyCG, the DNA-bending energy profile is invariant under a screw-like motion of DNA. Therefore, it is natural that this type of sequences will prefer to slide via a rotation-coupled motion that enables the nucleosome to maintain histone-DNA contacts at most times. This situation changes after the introduction of nucleosome positioning elements such as the TA motifs found in the polyCG-601 and 601 sequences considered here. TA motifs are highly flexible and they prefer to localize at the histone-DNA contact points where the DNA minor groove bends inward towards the histone core [8]. Starting from the initial optimal configuration, a DNA screw-like motion by 5 base pairs would bring these positioning motifs to the locations where the minor groove bends outwards, generating an unfavorable DNA bending profile. When many of these motifs are present, as in the 601 case, the bending energy penalty due to DNA rotation may become high, and repositioning proceeds instead via the observed rotation-uncoupled route. In this way, the optimal DNA bending profile is preserved at the expenses of hydrogen bonds, which are broken to form an intermediate 5-bp-shifted state. Hydrogen bonds are then restored after a further 5-bp jump. Notably, in the rotation-uncoupled sliding of the 601 sequence, we did not find reptation-like movement via a loop defect.

In our MD simulations, to keep the model simple and to enhance the observation of repositioning events, we employed the smallest hydrogen bond strength that can stabilize the nucleosome to the observed experimental crystal structure. Using these settings, we obtained results consistent with the experimental nucleosome unwrapping profiles as a function of salt concentration. However, fine tuning of the relative strength of nucleosome interactions at this level of coarse-graining represents a very challenging problem and we should consider the possibility that stronger hydrogen bond interactions may be more appropriate to represent the behavior of the system. Therefore, we reweighted the observed repositioning free energy profiles using higher hydrogen bonds strengths, and deduced how the two repositioning modes will be affected. Hydrogen bond strength has essentially no influence on the free energy profile of the rotation-coupled sliding observed in polyCG. On the other hand, for 601 we found that increasing the hydrogen bond strength by only 25% will increase the free energy difference between the optimal nucleosome conformation and the 5-bp-shifted state up to 10 *k*_B_*T,* a value comparable to the DNA-bending energy penalty that 601 nucleosomes should pay to reposition via the rotation-coupled mode [27]. Therefore, screw-like sliding may be the dominant mode even in the case of strong positioning sequences, as suggested by some authors [15].

While the current study successfully observed two distinct sliding modes via direct CGMD simulations, many related computational studies remain to be done for more comprehensive understanding. First, since the spontaneous sliding is inherently a rare event, one can utilize advanced sampling methods to enhance the observation of sliding events, such as transition path sampling [45] and Markov state modeling [46]. Related to it, one could reverse map the obtained CG structural ensemble of sliding intermediates to fully atomistic models, as was done in Ref. [47], so that the dynamics of nucleosome sliding could be analyzed at atomic resolution. Furthermore, while the current simulations assume a stable histone octamer, the model could be improved by including histone octamer disassembly [39], which could be of particular importance to study nucleosome distortions induced by the action of active chromatin remodelers [10]. Finally, it would be interesting to apply our model to study the effect of sliding on the dynamics of large-scale chromatin fibers [48,49].

## Conclusion

In summary, we firstly developed a novel representation of protein-DNA hydrogen bonds at the coarse-grained level and confirmed their importance for nucleosome stability. Then, we investigated the kinetics of nucleosome sliding for 3 sequences and found 2 distinct repositioning modes: rotation-coupled and rotation-uncoupled. The sequence-dependent intrinsic flexibility of DNA determines not only nucleosome position, but also the kinetics of nucleosome sliding. The underlying mechanism to switch the sliding modes is a balance between hydrogen bond interaction and DNA bending energy. For non-positioning sequences, screw-like rotation of the DNA always represents the dominant repositioning strategy. However, when DNA has a strong intrinsic bending due to the presence of positioning motifs, the bending energy penalty due to DNA rotation may become too large, and a different rotation-uncoupled mode proceeding via the breakage of all histone-DNA hydrogen bonds may dominate. For both sliding modes, due to the free energy barriers of DNA bending or hydrogen bond breakage, spontaneous sliding of positioning sequences is much slower than that of uniform sequences, suggesting that the action of active chromatin remodelers may be required only when the nucleosome sequence is rich in positioning motifs such as TA base pair steps.

